# Regulation of Epithelial HIF by Probiotic *Escherichia coli*

**DOI:** 10.1101/2025.05.14.654147

**Authors:** Alexander S. Dowdell, Rebecca Roer, Geetha Bhagavatula, Ian M. Cartwright, Rachel H. Cohen, Jacob A. Countess, Samuel D. Koch, J. Scott Lee, Calen A. Steiner, Noah T. Thompson, Zachary F. Villamaria, Nichole M. Welch, Corey S. Worledge, Liheng Zhou, Sean P. Colgan

**Affiliations:** Mucosal Inflammation Program and Division of Gastroenterology and Hepatology, Department of Medicine, University of Colorado School of Medicine, Aurora, CO, USA; Rocky Mountain Regional Veterans Affairs Medical Center, Aurora, CO, USA; Department of Pediatrics, Children’s Hospital Colorado, University of Colorado, Aurora, Colorado

## Abstract

The gastrointestinal tract is home to trillions of microorganisms that interact with their host in profound ways, including regulation of immune, endocrine, and neurological functions. One mechanism by which these microbes interact with their eukaryotic host is through the generation of short-chain fatty acids (SCFAs), which are metabolized by the intestinal epithelium creating a state of “physiologic hypoxia”. This hypoxia, in turn, results in stabilization and activation of hypoxia-inducible factor (HIF), a transcription factor family shown to support gut barrier function and homeostasis, in the intestinal epithelium. The association between HIF and intestinal homeostasis has been long understood, as both genetic and pharmacologic potentiation of the HIF signaling pathway has been shown to promote barrier function both *in vitro* and *in vivo*. Although it has been previously established that pathogenic bacteria regulate HIF stabilization and activity in the intestinal epithelium independent of SCFA metabolism, it is not clear whether this property extends to noninfectious and/or commensal bacterial species. Here, we demonstrate that nonpathogenic, commensal strains of *Escherichia coli* stabilize HIF in intestinal epithelial cells *in vitro*. Further, we show that HIF is transcriptionally active in these cells and drives a “pro-barrier” transcriptional program. This property was found to be dependent on bacterial aerobic respiration, as genetic elimination of *E. coli* aerobic respiration abolished HIF stabilization and the subsequent transcriptional phenotype. Finally, we observed induction of tissue hypoxia *in vivo* using antibiotic-treated mice colonized with wild-type, but not respiration-deficient, *E. coli.* These findings demonstrate a novel ability for probiotic *E. coli* to regulate intestinal homeostasis through activation of HIF and suggest that this mechanism might be leveraged in as a novel therapeutic to combat intestinal inflammation, such as that observed during inflammatory bowel disease (IBD).

## Introduction

Inflammatory bowel disease (IBD) is a family of conditions characterized by chronic, relapsing inflammation of the gastrointestinal tract^1^. IBD can be divided into two general sub-types (Crohn’s disease and ulcerative colitis) that share some similarities but diverge in their specific pathology and genetic risk factors^2^. The pathogenesis of IBD is a complex interplay of genetic, environmental and microbial factors; as a result, a full understanding of the pathophysiological mechanisms that drive disease in the various IBD sub-types has not been achieved^3^. Currently, no cure for IBD exists and current treatments are plagued by complications such as lack of response/loss of efficacy, immunosuppression, and risk of infection/cancer^4^. As a result, there exists a present need for a better understanding both of the healthy gut physiology as well as the pathological mechanisms of IBD in order to inform the development of novel treatments.

One striking feature of IBD is the impairment of intestinal “barrier function”, that is, the gut’s ability to selectively absorb nutrients and water while excluding microbes and toxins^5^. This loss in selective permeability, termed “leaky gut”, permits the translocation of microbes and microbially derived molecules (such as lipopolysaccharide), resulting in systemic inflammation^6^. Interestingly, increased gut permeability has been found to accurately predict future Crohn’s disease in asymptomatic, first-degree relatives of Crohn’s disease patients^7^. Similarly, changes in gut barrier functions have been found to be a better predictor of disease prognosis than either histologic or endoscopic remission for both ulcerative colitis and Crohn’s disease^8^. These findings underscore the importance of normal barrier function in intestinal homeostasis and suggest that potentiation of gut barrier function may be a viable strategy for the treatment of IBD. Intestinal barrier function is regulated by numerous molecular and environmental factors, including dietary, medicinal, and genetic influences^9^. One such cell-intrinsic component to intestinal barrier function is the transcription factor family hypoxia-inducible factor (HIF)^10^. The HIF transcriptional unit is composed of an oxygen-labile α-subunit, of which three paralogs have been identified (HIF-1α/-2α/-3α), and an oxygen-insensitive β-subunit (HIF-1β/ARNT)^11^. The stability of HIF-α proteins is regulated post-translationally by a family of prolyl hydroxylases (PHDs) which, under conditions of replete oxygen, ferrous iron, ascorbate, and α-ketoglutarate, hydroxylate HIF-α at particular proline residues as a marker for poly-ubiquitination and proteasomal degradation^12^. During cirumstances in which any of these substrates becomes limiting (most notably, during hypoxia), the hydroxylation activity of PHDs becomes compromised and, as a result, HIF-α subunits accumulate in the cytoplasm. From there, these subunits dimerize with HIF-1β/ARNT, translocate to the nucleus, and exert specific transcriptional effects that can vary by cell and tissue^11^. HIF signaling is active at homeostasis in the intestinal epithelium due to the tissue’s “physiologic hypoxia”, a state driven by a combination of factors including “counter-current” gas exchange between neighboring venous and arterial capillaries in intestinal villi and by epithelial metabolism of bacterially derived short-chain fatty acids (SCFAs) through β-oxidation^13^. Intestinal HIF expression has been shown to drive a number of “pro-barrier” factors including intestinal trefoil factor (ITF/TFF3), claudin-1, and mucin-3^14^. Mice deficient in intestinal HIF-1α expression, correspondingly, show increased metrics of disease in mouse models of IBD, while mice treated with compounds that stabilize HIF through inhibition of PHDs are conversely protected in such models^15–17^. These findings demonstrate that HIF signaling, through multiple mechanisms, promotes gut homeostasis and is protective during intestinal inflammation.

HIF signaling has been shown to be modulated by extrinsic factors, in addition to the intrinsic regulation by PHDs. Previously, our group and others demonstrated that pathogenic bacteria stabilize HIF-1α in epithelial cells and activate HIF-1α-dependent transcriptional pathways^18^. This phenomenon was found to be true for other, diverse pathogenic species, including *Staphylococcus aureus*, *Yersinia enterocolitica*, *Borrelia burgdorferi*, and *Acinetobacter baumannii*^18, 19^. Our group subsequently found that HIF stabilization induced by the model invasive bacterium *Salmonella enterica* subsp. *enterica* serovar Typhimurium (hereafter, “*Salmonella* Typhimurium”) activated anti-bacterial autophagy (“xenophagy”) in intestinal epithelial cells (IECs) in a HIF-dependent manner^20^. These findings provided a link between HIF stabilization, autophagy, and gut homeostasis; however, as our results indicated an oxygen-dependent mechanism for HIF-1α stabilization, we speculated that bacterial invasion might be dispensable for HIF-1α stabilization in IECs. In this work, we show that noninvasive *Salmonella* Typhimurium stabilizes HIF-1α to the same extent as wild-type bacteria. Further, we demonstrate that probiotic, commensal *Escherichia coli* strains stabilize HIF-1α and activate HIF transcriptional programs in epithelial cells. Finally, we recapitulate our *in vivo* findings using antibiotic-treated mice, demonstrating that wild-type, but not respiration-deficient, *E. coli* induces tissue hypoxia in the cecum. Taken together, these findings demonstrate a novel role for commensal bacterium such as *E. coli* in the regulation of intestinal homeostasis through activation of HIF signaling pathways.

## Materials and Methods

### Cell Lines and Bacterial Cultures

HeLa cervical adenocarcinoma cells (ATCC #CCL-2) and C2BBe1 colorectal adenocarcinoma cells (ATCC #CRL-2102) were grown as described previously^20–22^. Briefly, cells were grown in Iscove’s Modification of DMEM (IMDM, Corning #10-016-CV) supplemented with 10% (v/v) heat-inactivated bovine calf serum (BCS, Cytiva #SH30072.03) and 1x GlutaMAX (Thermo #35050061). hIEC-6 cells (ATCC #CRL-3266) were grown in Opti-MEM (Thermo #31985-070) supplemented with 20 mM HEPES (Thermo #15630-080), 10 mM GlutaMAX, 10 ng/mL epidermal growth factor (Corning #354052), and 4% (v/v) fetal bovine serum (ATCC #30-2021). All cells were grown at 37 °C, 5% CO_2_ in a humidified incubator. For experiments involving bacterial-cell co-cultures, cells were plated in 6-well tissue culture-treated plates at approximately 2.4 x 10^5^ cells/well the day before the experiment. C2BBe1 cells were plated onto collagen-coated 6-well plates (Corning #354400) to improve adherence.

Bacterial strains used in these experiments are listed in **Table 1**. Bacteria were routinely cultured in LB-Miller broth (BD #244620) at 37 °C, 250 RPM and maintained on LB-Miller plates solidified with 1.5% agar (BD #214010). Plates were freshly struck on a weekly or a biweekly basis from 15% (v/v) glycerol stocks kept at -80 °C. Antibiotics were included at the following working concentrations, as needed: 100 μg/mL carbenicillin (Sigma-Aldrich #C1389), 100 μg/mL streptomycin (Sigma-Aldrich #S6501), 100 μg/mL spectinomycin (Sigma-Aldrich #S0692), 100 μg/mL kanamycin (Sigma-Aldrich #60615), and 25 μg/mL chloramphenicol.

**Table 1.**
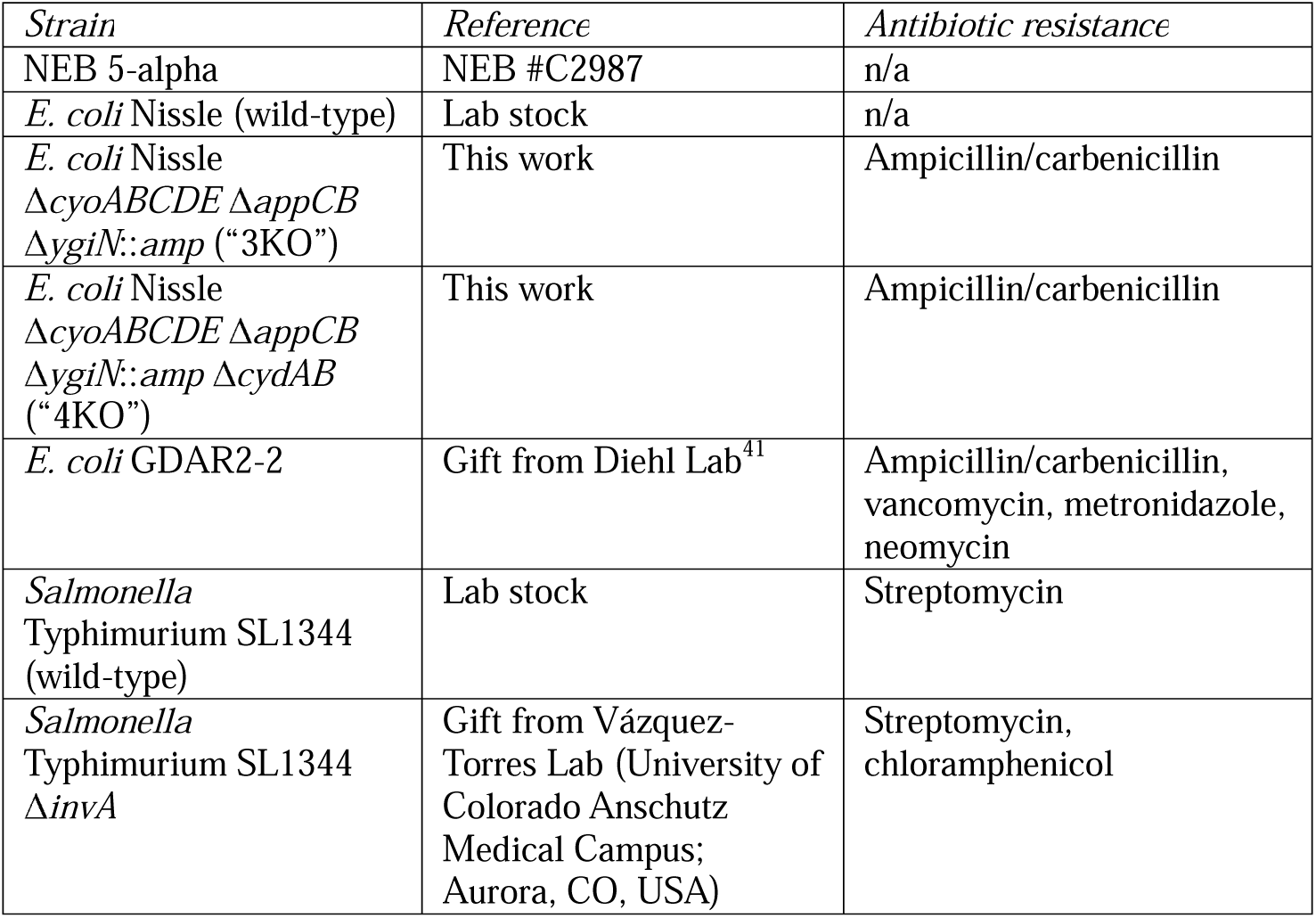
Bacterial strains used in this study.

| *Strain* | *Reference* | *Antibiotic resistance* |
| --- | --- | --- |
| NEB 5-alpha | NEB #C2987 | n/a |
| *E. coli* Nissle (wild-type) | Lab stock | n/a |
| *E. coli* Nissle<br>ΔcyoABCDE ΔappCB<br>ΔygiN::amp (“3KO”) | This work | Ampicillin/carbenicillin |
| *E. coli* Nissle<br>ΔcyoABCDE ΔappCB<br>ΔygiN::amp ΔcydAB<br>(“4KO”) | This work | Ampicillin/carbenicillin |
| *E. coli* GDAR2-2 | Gift from Diehl Lab<sup>41</sup> | Ampicillin/carbenicillin,<br>vancomycin, metronidazole,<br>neomycin |
| *Salmonella*<br>Typhimurium SL1344<br>(wild-type) | Lab stock | Streptomycin |
| *Salmonella*<br>Typhimurium SL1344<br>ΔinvA | Gift from Vázquez-Torres Lab (University of Colorado Anschutz Medical Campus; Aurora, CO, USA) | Streptomycin,<br>chloramphenicol |

Experiments involving the *E. coli* Nissle strain lacking all three cytochrome oxidases (*E. coli* Nissle “4KO”) were performed using BHI broth (BD #237500) and BHI agar (BD #211065). When experiments were performed that included *E. coli* Nissle 4KO, all other bacterial strains were similarly grown in BHI as a control. For cell treatments, bacterial overnight cultures were grown in 2 mL of medium with antibiotics, as necessary, using 14 mL loosely capped, round bottom tubes. 1 mL of overnight culture was then pelleted at 8,000 x *g* for 3 minutes at room temperature, with the resulting supernatant removed by pipetting. The bacterial pellet was resuspended in 1 mL of sterile PBS, pH 7.4 and used immediately for cell treatments.

### Construction of Mutant Strains

Genetically modified *E. coli* Nissle strains were constructed using CRISPR/Cas9 + λ-Red recombineering as described previously^23^. Strain creation was designed/planned *in silico* using SnapGene prior to *in vitro* experiments. Primer and guide RNA (gRNA) sequences used can be found in **Table 2**, with the latter given as the oligos needed for cloning into linearized pEcgRNA (see below). In brief, gRNA sequences were designed against the regions to be deleted (*cyoABCDE*, *appCB*, and *cydAB*) and validated against the *E. coli* Nissle genome using CHOPCHOP^24–26^. Three gRNA sequences (A, B, C) were designed and tested separately for each genomic locus to be edited. The plasmids pEcCas (Addgene #73227) and pEcgRNA (Addgene #166581) were gifts from Sheng Yang. Primers were designed to amplify the genomic regions immediately 5’ and 3’ of the regions to be deleted, with the 5’ reverse primer ending at the start codon of the first gene in the operon and the 3’ forward primer beginning at the stop codon of the last gene. These primers also contained overlaps to facilitate covalent linkage of these fragments for the creation of recombineering donor DNA. The result is that the donor DNA for λ-Red recombineering contains an in-frame deletion of the coding DNA sequence of each of the deleted genes, minimizing the likelihood of polar effects on nearby genes. PCR fragments from the 5’ and 3’ regions were amplified using high-fidelity polymerase (Q5 Hot Start, NEB) and gel-purified. Fragments were then covalently joined using NEBuilder HiFi Master Mix to reduce the chance of annealing errors; the resulting covalently linked fragment was then PCR amplified and gel-purified. pEcgRNA was linearized by digestion with BsaI-HFv2 (NEB), and the correctly sized fragment was gel-purified. Linearized pEcgRNA and gRNA oligos were ligated as described elsewhere and cloned into chemically competent NEB 5-alpha cells (NEB). Plasmids were purified from the resulting transformants and sequenced using Nanopore whole plasmid sequencing (Quintara Biosciences). *E. coli* Nissle was made electrocompetent as described elsewhere using ice-cold 10% glycerol and stored as 30 μL aliquots at -80 °C^27^. Electrocompetent *E. coli* Nissle was first transformed with pEcCas, then subsequent electrocompetent cells were prepared with the inclusion of 10 mM L-arabinose in sub-cultures in order to induce expression of λ-Red genes. Approximately 30 ng of pEcgRNA containing the appropriate gRNA was electroporated alongside about 120 ng of full-length donor DNA to delete each operon of interest. Electroporations were performed using a Gene Pulser Xcell Electroporation System with 0.2 cm cuvettes, using the pre-programmed exponential decay setting “Bacteria -> *E. coli* -> 2.5 kV, 0.2 cm”. Putative knockouts were screened by PCR, with the absence of any other mutations confirmed by Sanger sequencing of the manipulated loci (ACGT, Inc.). After confirmation of the success of each mutation, the pEcgRNA plasmid was cured using 10 mM L-rhamnose induction to allow for subsequent mutations to be made.

**Table 2.**
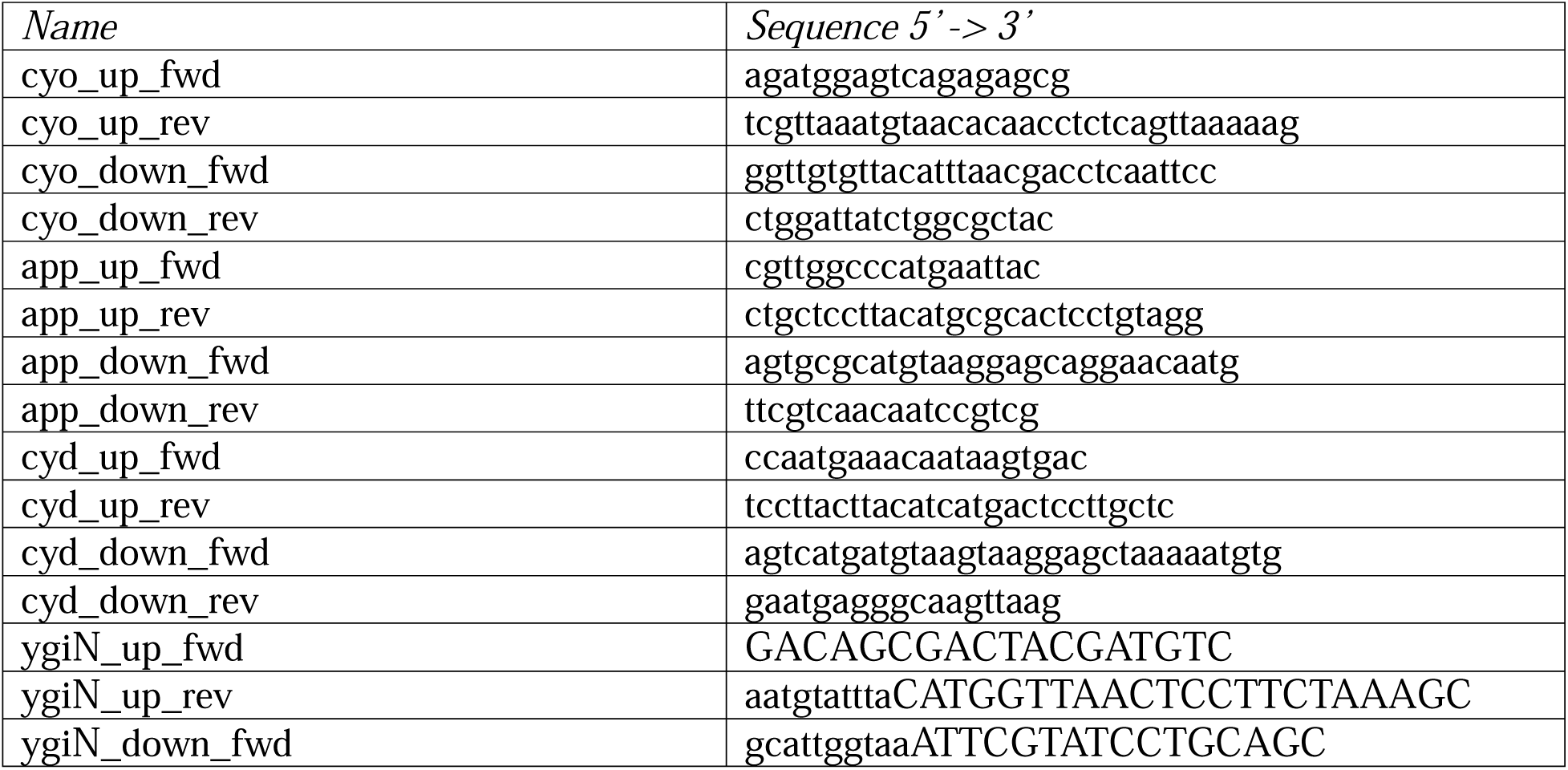
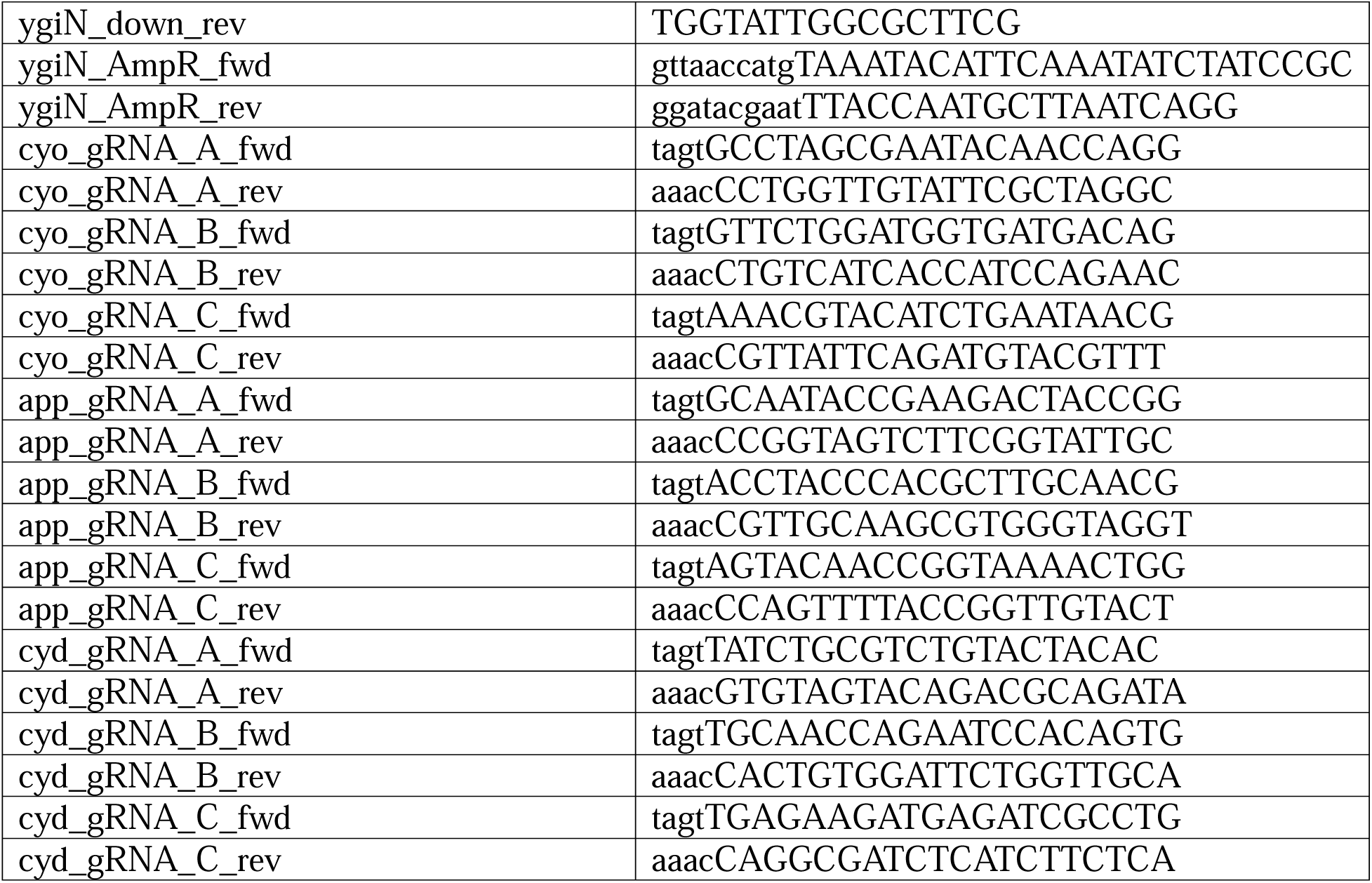
Primers and oligos used for construction of mutant strains.

Deletion of *ygiN* occurred as for the others except that an ampicillin resistance cassette was included as a third fragment in between the 5’ and 3’ fragments, with the stop codon of the *ampR* gene occurring in-frame and eight codons prior to the *ygiN* stop codon. The ampicillin resistance cassette was amplified from pM1s3AsG (Addgene #137921, a gift from Neel Joshi) using the given primers and covalently joined using NEBuilder HiFi Master Mix as for the other donor DNA fragments^28^. As the presence of the ampicillin resistance cassette provided a suitable means of counter-selection for wild-type *ygiN* cells, no gRNA was utilized for this knockout.

Mutations were made in the order Δ*cyoABCDE ->*Δ*appCB* -> Δ*ygiN*::*amp* -> Δ*cydAB*. Following knockout of both Δ*cyoABCDE* and Δ*cydAB*, a growth defect in LB-Miller medium was observed as previously reported ^29^. Growth of these strains was observed to be substantially more robust on a more enriched medium; hence, BHI was used for the routine propagation of these strains.

### Cell Treatments and Immunoblotting

Analysis of HIF-1α by immunoblotting was performed as described previously^20, 30^. Briefly, cells plated in 6-well plates were fed with fresh cell culture medium and treated for the given time period, typically 6 hours, with the given strain at a standardized inoculum of 4 x 10^7^ colony forming units (CFU) per well or with 300 μM cobalt chloride as a positive control for HIF-1α stabilization. In some cases, bacteria were killed by heating at 65 °C or by treating with glutaraldehyde at 5% (w/v) final concentration for approx. 2 hrs. 15 min. each. Confirmation of killing efficacy was confirmed by plating undiluted, treated cells onto LB-Miller agar and incubating overnight at 37 °C – in either case, the absence of colonies was observed indicating sterilization of cultures. Prior to use, glutaraldehyde-treated cells were thoroughly washed with PBS to remove residual glutaraldehyde. In other cases, 0.4 μm pore size cell culture inserts inserts (Greiner #657640) were placed above the cells, with 1 mL of medium added to the inner “apical” chamber and with/without the same inoculum of EcN that wells without inserts received. After the given time, plates were placed on ice and medium aspirated. Cells were then immediately lysed using ice-cold lysis buffer, consisting of 1x Laemmli sample buffer (Bio-Rad #1610747), 100 mM freshly made DTT (Sigma-Aldrich #D0632), 1x HALT Protease Inhibitor Cocktail (Thermo #78438), and 0.5 mM EDTA. Lysates were transferred to 1.5 mL microcentrifuge tubes and sonicated to reduce viscosity, then frozen at -20 °C until needed for analysis. Lysates were separated by SDS-PAGE using 4-20% gradient pre-cast gels (Bio-Rad #4568094 or 4568096), then transferred to 0.2 μm PVDF using a Bio-Rad Trans Blot Turbo instrument and RTA Mini Kits (Bio-Rad #1704272). Subsequent blots were blocked for one hour at room temp. with 5% (w/v) nonfat dry milk (Bio-Rad #1706404) in tris-buffered saline + 0.1% (v/v) Tween 20 (TBS-T) (“blocking buffer”) and incubated with a primary antibody overnight at 4 °C. Primary antibodies used in this study were mouse anti-HIF1A (BD #610959, 1/500) and rabbit anti-ACTB (Abcam #ab8227, 1/10,000) and were diluted in blocking buffer. Blots were then washed repeatedly with TBS-T and incubated with HRP-conjugated secondary antibody (MP Bio. #0855676, 0855550) diluted 1/10,000 in blocking buffer for one hour at room temp. Blots were then further washed with TBS-T and developed using Clarity ECL reagent (Bio-Rad #1705061), then imaged using a Bio-Rad ChemiDoc MP instrument.

### RNA Purification, Quantitative PCR, and Gene Expression Analysis

Cells were treated similarly as for immunoblotting except that treatment time was extended to 8 hours and inoculum was approx. 10^7^ CFU/well. At the end of the treatment time, medium was aspirated and 1 mL of cold TRIzol reagent (Thermo #15596026) was added to each well. Plates were swirled gently to lyse cells, then lysates were transferred to microcentrifuge tubes and frozen at -80 °C. RNA was purified from TRIzol lysates according to manufacturer’s instructions except that 1-bromo-3-chloropropane was used as the phase separation reagent^31^.

Purified RNA was then cleaned up using lithium chloride (Sigma-Aldrich #L7026) precipitation according to established protocols (product info. sheet for Thermo #AM9480). cDNA was then synthesized from RNA using iScript Supermix (Bio-Rad #1708840). Gene expression was measured using Power SYBR Green PCR Master Mix (Thermo #4367659) and with a QuantStudio 3 Real-Time PCR System, using the qPCR primers given in **Table 3**. Removal of residual gDNA by LiCl cleanup was confirmed using no-RT Control Supermix included with iScript Kit. Primer sequences were obtained from Harvard PrimerBank, with PrimerBank ID numbers listed in **Table 3**^32^. Gene expression analysis was conducted using LinRegPCR^33^.

**Table 3.**
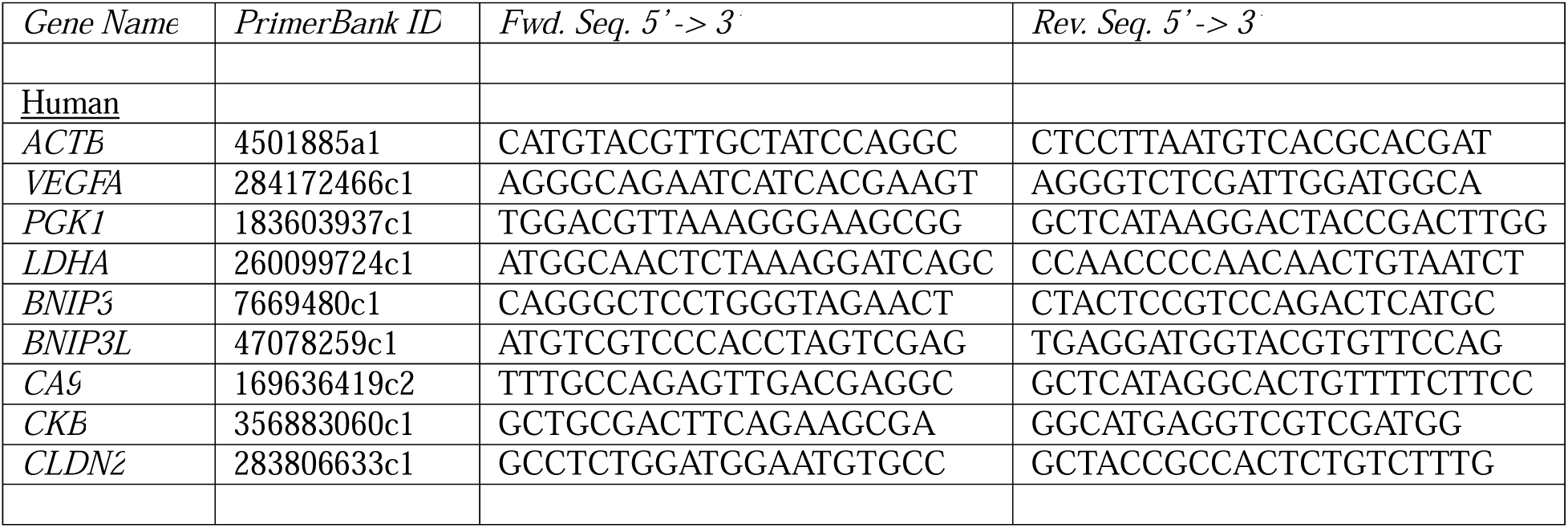
Quantitative PCR (qPCR) primers used in this study.

Results are representative of at least two independent experimental repetitions.

### Oxygen Consumption Experiments

Oxygen consumption experiments were performed using OxoDish OD24 plates (PreSens Precision Sensing GmbH), which measure dissolved oxygen level through a precalibrated fluorescent probe. Overnight cultures of bacterial strains were prepared as described above, then inoculated into DMEM/F12 (Thermo #1133032) after normalization based on OD600 and incubated with gentle rocking at 37 °C. Some wells contained only media as a negative control. Bacteria for inoculation were prepared as described before for cell treatments. Experiment was performed at least twice with similar results in each repetition.

### Luciferase Reporter Experiments

HeLa cells were plated on 6-well plates, then co-transfected approximately one day later with pGL4.22-PGK1-HRE::dLUC (Addgene #128095, a gift from Chi Van Dang), which expresses destabilized firefly luciferase from a HIF-1α-responsive promoter, and pGL4.75[hRluc/CMV] (Promega #E6931) as a normalization control, which constitutively expresses *Renilla* luciferase from a CMV promoter^34^. Transfections were performed using FuGENE HD (Promega #E2311) according to the manufacturer’s instructions. One day after transfection, cells were treated in fresh growth medium with either 300 μM CoCl_2_ (positive control) or with about 2.7 x 10^6^ CFU/well of the indicated strain of *E. coli* Nissle. Some wells were left untreated as negative controls. Cells were incubated with treatments for 8 hours at 37 °C, 5% CO_2_, then luciferase activity was measured using a Dual-Glo Luciferase Assay System (Promega #E2920) according to the manufacturer’s protocol. Luciferase measurements were taken using a Biotek Synergy H1 plate reader. Results are representative of at least two independent experimental repetitions.

### Salmonella Invasion Assays

Invasiveness of wild-type *Salmonella* Typhimurium SL1344 and the otherwise isogenic mutant SL1344 Δ*invA* were measured as described in detail elsewhere^35^. Briefly, HeLa cells were infected with *Salmonella* Typhimurium at a multiplicity of infection (MOI) of approx. 151 (SL1344 WT) or 217 (SL1344 Δ*invA*) for 10 min., after which cells were washed and given fresh antibiotic-free medium. At approx. 30 min. post-infection, gentamicin (Lonza #17-518Z) was added to cells at 100 μg/mL to kill extracellular bacteria. At 1 hour post-infection, medium was aspirated from cells, and cells were thoroughly washed with sterile PBS. Cells were then lysed using PBS + 0.2% (w/v) sodium deoxycholate (Sigma-Aldrich #D6750). Lysates were serially diluted using sterile PBS and plated onto LB-Miller agar containing 100 μg/mL streptomycin.

Plates were incubated overnight at 30 °C. Colonies were counted the following day, and colony counts used to determine number of intracellular bacteria.

### In vivo colonization and hypoxia assessment

Wild-type C57BL/6 mice were maintained at the University of Colorado Anschutz Medical Campus in an AAALAC-accredited (#00235) and USDA-inspected (#84-R-0059) facility. All animal procedures were performed under an approved IACUC protocol (#00182). Broad-spectrum antibiotic treatments were performed as described previously^36^. In brief, mice were gavaged with 100 μL of a sterile-filtered antibiotic solution containing 2 mg/mL ampicillin (Sigma-Aldrich #A0166), 2 mg/mL gentamicin (Sigma-Aldrich #G1914), 2 mg/mL metronidazole (Sigma-Aldrich #M1547), 2 mg/mL neomycin (Sigma-Aldrich #N6386), and 1 mg/mL vancomycin (Sigma-Aldrich #V2002) in Milli-Q water. Control mice received no antibiotics. Gavages were performed for three consecutive days, after which mice were given a 24-hour period to allow “wash out” of antibiotics from the gastrointestinal tract. After this, mice were gavaged with ∼10^8^ – 10^9^ CFU of *E. coli* Nissle wild-type or “4KO” respiration-deficient mutant in 100 μL sterile PBS. Both strains contained the pEcCas plasmid as a means of conferring kanamycin resistance – this allowed for accurate determination of bacterial density from fecal pellets. Presence of this plasmid had no observable effects on bacterial growth. *E. coli* was grown and prepared in PBS as indicated above. As a control, some mice received no *E. coli* at this. Mice received two days of *E. coli* gavages, after which each mouse (including controls) was intraperitoneally injected with 15 μg/μL of pimonidazole (HP-200mg, Hypoxyprobe Inc.) freshly suspended in sterile PBS. After 60 minutes, mice were sacrificed by CO_2_ asphyxiation + cervical dislocation, and cecal tissue was isolated into 10% neutral buffered formalin for histology. The experiment was repeated twice with representative tissue images given. Colonization of bacteria was determined by plating fecal pellets obtained just prior to sacrifice: fecal pellets were homogenized in sterile PBS and plated onto either LB-Miller agar (EcN WT) + 100 μg/mL kanamycin or BHI + 100 μg/mL ampicillin and kanamycin. In all mice, colonization after 2 days was approximately 4 x 10^7^ – 1 x 10^8^ CFU/g feces regardless of EcN strain used.

### Immunohistochemistry (IHC) detection of pimonidazole (Hypoxyprobe) in tissues

Murine cecal tissue was fixed in 10% neutral buffered formalin immediately following euthanasia and necropsy. Fixed cecal tissues were stored at 4 °C until processed through embedding in paraffin. Tissue sections were cut and stained for pimonidazole using rabbit anti-pimonidazole antibody (Pab2627, HypoxyProbe Inc.) using established techniques^70^. Peroxidase-developed tissue sections were counterstained with hematoxylin and photographed using a Zeiss AxioImager A1 microscope + AxioCam MRc 5 camera.

### Data Analysis, Statistical Testing, and Image Generation

Data were analyzed using GraphPad Prism 10 or Microsoft Excel. Statistical tests used can be found in the figure legend for the appropriate figure. Figure images were prepared using GraphPad Prism, Microsoft PowerPoint, SnapGene, or BioRender.

## Results

Previously, our group sought to characterize the role of HIF-1α in the regulation of anti-bacterial autophagy (“xenophagy”) occurring during infection of epithelial cells by *Samonella* Typhimurium (STm)^20^. Our group found that HIF-1α positively regulated this process, and that HIF-1α was stabilized in the course of infection by STm. During these studies, we found that treatment of intestinal epithelial cells *in vitro* with STm markedly reduced oxygen tensions, a potential mechanism for the observed stabilization of HIF-1α. As STm is a facultative anaerobe and potentially could be respiring oxygen extracellularly, we sought to clarify whether invasion of epithelial cells is a prerequisite for HIF-1α stabilization in a bacterial/epithelial co-culture model. To answer this question, we treated epithelial cells with either wild-type STm SL1344, a highly invasive strain of *Samonella*, or an otherwise isogenic mutant deficient in expression of the type 3 secretion system protein InvA (SL1344 Δ*invA*). *Salmonella* Typhimurium expresses two pathogenicity islands necessary for invasion of eukaryotic cells and replication therein; disruption of certain crucial genes in the first island, such as InvA or InvG, has been previously shown to abrogate the ability of STm to invade epithelial cells^37–39^. Measurement of the invasiveness of these strains in HeLa cells shows that wild-type STm SL1344 invade cells aggressively, whereas invasion is essentially absent in the SL1344 Δ*invA* mutant (**Figure 1a**). Next, we treated HeLa cells with each mutant at varying MOIs or for varying times. As indicated in **Figure 1b** and **Figure 1c**, the wild-type strain robustly stabilized HIF-1α in a time- and dose-dependent fashion (respectively) as previously observed^20^. Surprisingly, the mutant Δ*invA* strain stabilized HIF-1α to an equal extent as the wild-type strain, indicating that cellular invasion is not a prerequisite for HIF-1α stabilization in epithelial cells.

**Figure 1:**
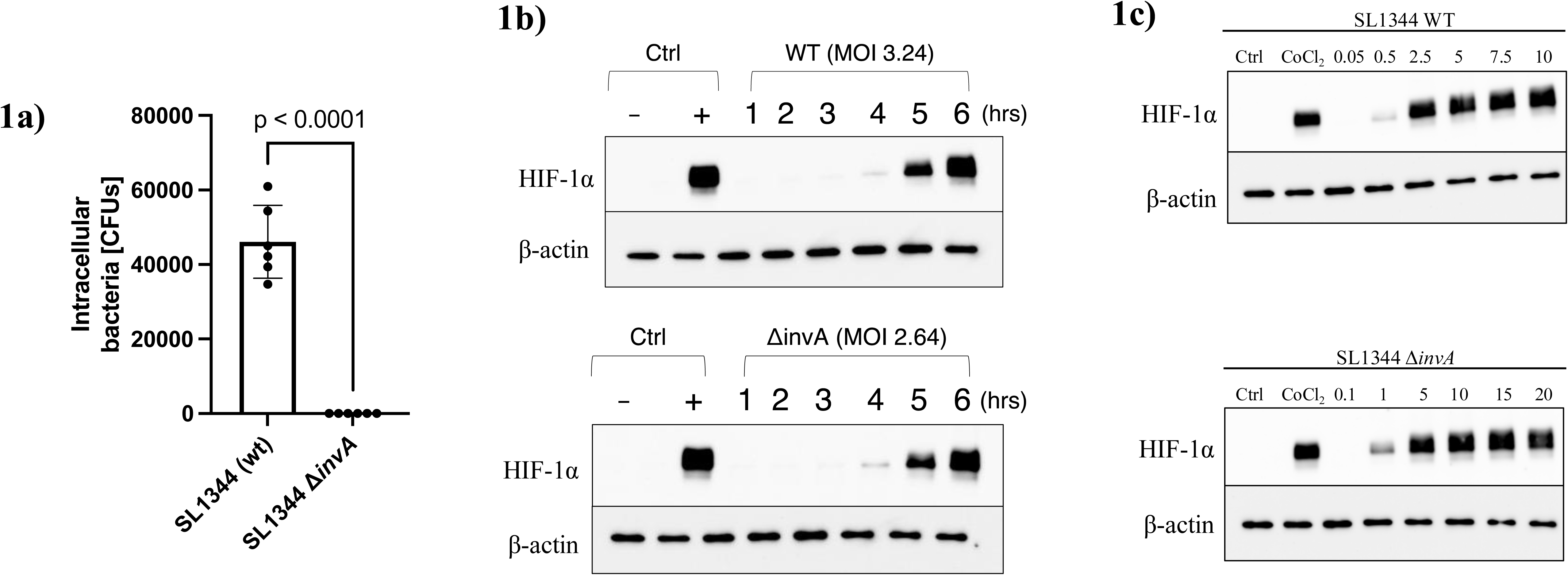
Invasion is not a prerequisite for HIF-1α stabilization by *Salmonella* Typhimurium (STm). a) Invasion assay in HeLa cells with either STm SL1344 wild-type (WT) or InvA-deficient mutant. Statistical significance calculated using an unpaired t test with Welch’s correction. Error bars represent standard deviation. (b) Western blot of HeLa cells treated at the indicated multiplicity of infection (MOI, initial bacterial CFU per cell) for varying times. Negative controls are untreated cells, while positive controls are cells treated with 300 μM CoCl_2_ for 6 hours. (c) Dose response of HeLa cells treated with SL1344 WT or InvA-deficient mutant. Cells treated at the indicated MOI for 6 hours then lysed. “Ctrl” = negative, untreated control cells. “CoCl_2_” = 300 μM CoCl_2_ for 6 hours.

As STm mutants deficient in cellular invasion have been previously shown attenuated pathogenicity *in vivo*, we then asked whether closely related, nonpathogenic bacteria might also stabilize HIF-1α *in vitro*^40^. To this end, we examined the ability of *Escherichia coli* to stabilize HIF-1α using either commensal/probiotic strains isolated from either human (*E. coli* Nissle, EcN) or mouse (GDAR2-2) sources^41, 42^. We also sought to answer whether our earlier findings could be reproduced in an intestinal epithelial cell line, such as the Caco-2 derivative C2BBe1. As can be seen in **Figure 2a**, both *E. coli* strains demonstrate robust HIF-1α stabilization indicating that pathogenicity in and of itself does not drive HIF-1α stabilization. In addition, we observed robust HIF-1α stabilization in C2BBe1 cells indicating that our previous findings were not specific to HeLa cells. We further asked whether the stabilized HIF-1α is transcriptionally active: HIF-1α is subject to direct post-translational regulation through modifications such as asparagine hydroxylation by FIH and indirect regulation such as cullin neddylation that regulate HIF-1α’s ability to affect cellular gene expression^43, 44^. To this end, we treated C2BBe1 cells with cobalt chloride (CoCl_2_), a known HIF-1α stabilizing agent, or EcN^45^. As shown in **Figure 2b**, treatment with either CoCl_2_ or EcN induces the expression of classic HIF-1α target genes such as *CA9*, *BNIP3*, *BNIP3L/NIX*, *LDHA, PGK1* (EcN only), and *VEGFA*^46–50^. Further, EcN significantly induced the expression of the “pro-barrier” target *CKB* while decreasing expression of *CLDN2*, whose expression has been shown to inversely correlate with barrier function^22, 51^. These findings suggest a possible protective role for EcN in maintaining intestinal epithelial barrier function through activation of HIF-1α.

**Figure 2:**
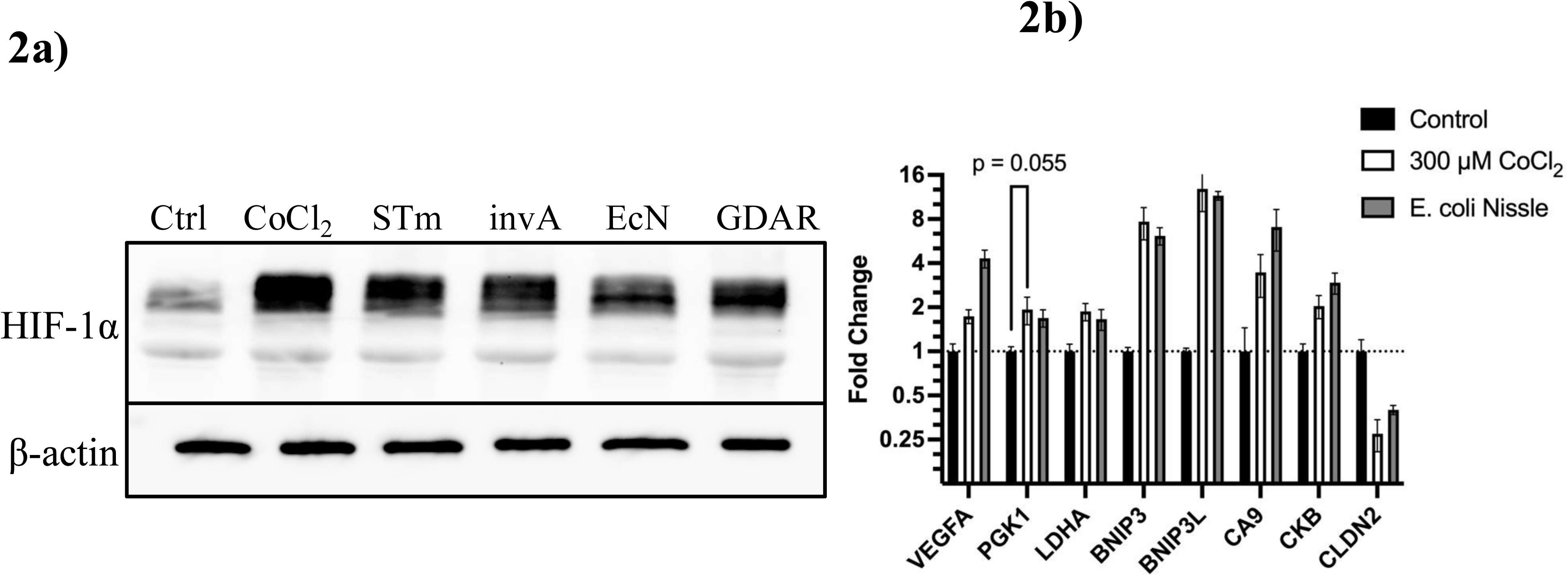
HIF-1α is stabilized by commensal *E. coli* and is transcriptionally active. (a) Western blot of C2BBe1 cells treated for 6 hours with approx. 4 x 10^7^ CFU/well of the indicated strains. “Ctrl” = untreated control cells, “CoCl_2_” = 300 μM CoCl_2_, “STm” = *Salmonella* Typhimurium SL1344 wild-type, “invA” = SL1344 Δ*invA*, “EcN” = *E. coli* Nissle, “GDAR” = *E. coli* GDAR2-2. (b) qPCR analysis of gene expression in C2BBe1 cells treated with either 300 μM CoCl_2_ or 10^7^ CFU/well *E. coli* Nissle for 8 hours. Data are normalized to β-actin expression and presented a fold-change relative to untreated control cells. Statistics calculated using unpaired t tests with Welch’s correction, comparing either CoCl_2_ treatment or EcN treatment samples to control (untreated) cells. All pairwise comparisons vs. control were significant (p < 0.05) except for PGK1 control vs. CoCl_2_, as indicated. Error bars represent standard deviation.

We next sought to elucidate the mechanism by which probiotic *E. coli* potentiates HIF-1α stability and transcriptional activity. We chose to focus on EcN, as this probiotic has received considerable recent attention as a potential chassis for detection and treatment of human gastrointestinal disease^52–54^. We asked whether live EcN is necessary for the stabilization of HIF-1α, as previous reports indicate substantial cross-talk between HIF-1α and NF-κB signaling pathways^55, 56^. Thus, we inactivated cultures of EcN using either moderate (65 °C) heating or with the crosslinking agent glutaraldehyde, which is used medically to sterilize surgical instruments ^57^. Inactivation of EcN using either heat or glutaraldehyde abolished bacteria-induced HIF-1α stabilization in treated HeLa cells (**Figure 3a**), suggesting that HIF-1α stabilization requires some biochemical process mediated by live bacteria. Further, we asked whether EcN might be secreting a soluble factor, such as lipopolysaccharide (LPS), responsible for the observed HIF-1α stabilization as LPS has been previously shown to activate HIF-1α in macrophages^58^. To answer this question, we added cell culture inserts above the plated HeLa cells just prior to treatment – these inserts have pore sizes of 0.4 μm which allow the passive diffusion of small, soluble factors but restrict the passage of *E. coli* (**Figure 3b**). Addition of a medium-containing insert had no effect by itself on HIF-1α stabilization (**Figure 3a**).

**Figure 3:**
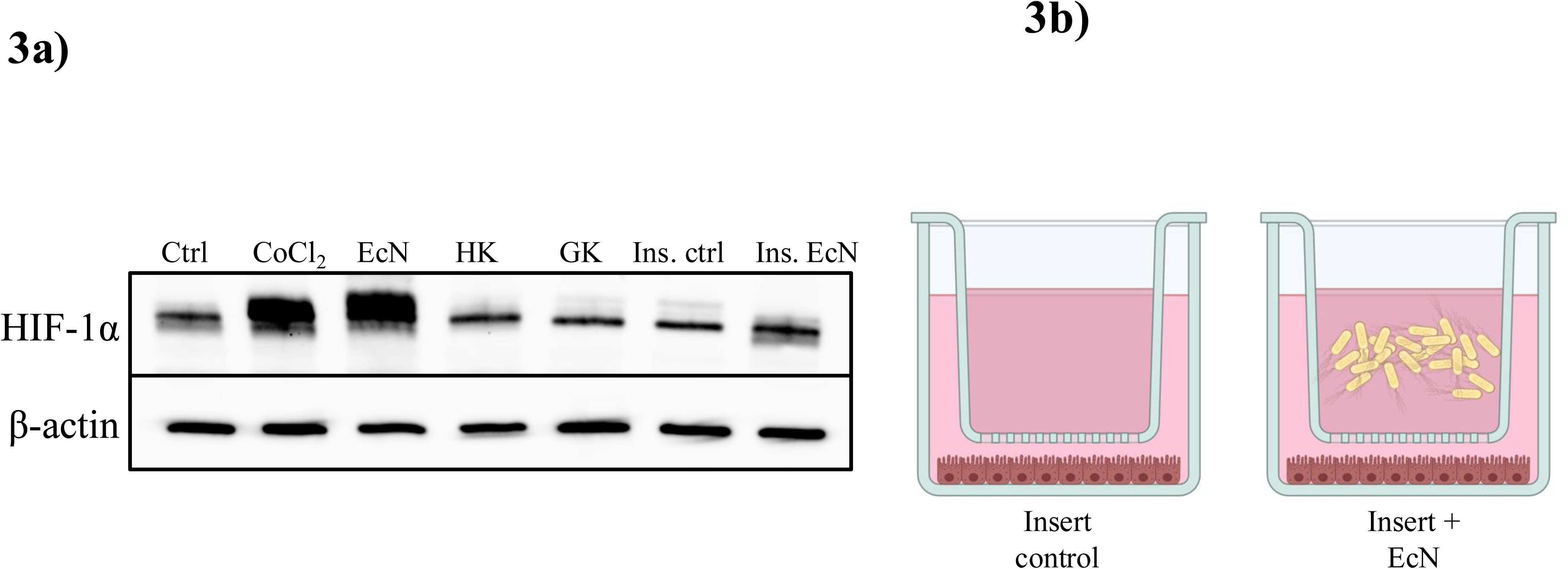
HIF-1α stabilization phenotype requires live, cell-associated bacteria. (a) HeLa cells treated as given for 6 hrs. at 37 °C, 5% CO_2_, then lysed for analysis by Western blotting. “Ctrl” = untreated (negative) control, “CoCl_2_” = 300 μM cobalt chloride (positive control), EcN = *E. coli* Nissle WT, “HK” = heat-killed EcN, “GK” = glutaraldehyde-killed EcN, “Ins. ctrl” = Cell culture insert, negative control, “Ins. EcN” = Cell culture insert containing EcN in inner, “apical” chamber. (b) Diagram of cell culture insert experiment. At left, control cells received only a steile insert so as to control for the effect of the insert on HIF-1α stabilization. At right, treated cells received 4 x 10^7^ CFU in the inner “apical” chamber of the insert, separated from the epithelial cells by a 0.4 μm pore size membrane that effectively blocked bacterial transmigration.

Interestingly, co-culture of EcN and HeLa cells separated by a permeable insert abrogated stabilization of HIF-1α (**Figure 3a**); these findings suggest that HIF-1α activation mediated by EcN is likely not due to a soluble factor and may require close cell-bacteria proximity.

Given our previous findings that live, cell-associated bacteria are required for stabilization of HIF-1α, we then asked what bacterial processes might drive HIF-1α activation in epithelial cells. As HIF-1α is stabilized classically by hypoxia, and as *E. coli* are facultative anaerobes, we hypothesized that HIF-1α stabilization in epithelial cells might be driven by bacterial aerobic respiration. As many of the classic electron transport chain inhibitors, such as oligomycin, antimycin A, and rotenone, are ineffective at fully blocking *E. coli* aerobic respiration, we turned to a genetic approach to abolish *E. coli* oxygen consumption^59–61^. Previous work suggests that deletion of *E. coli* cytochrome oxidases (Cyo, Cyd, and App/Cbd/Cyx), the central enzyme complexes that reduce molecular oxygen to water in the bacterial inner membrane, eliminates aerobic respiration and results in a fermentative growth phenotype^29, 62, 63^. To this end, we generated a strain of EcN which lacks all three cytochrome oxidases (Δ*cyo*ABCDE Δ*appCB* Δ*cydAB*) through a combination of λ-Red recombineering + CRISPR/Cas9 negative selection for bacteria that fail to undergo homologous recombination at the appropriate locus. A before/after summary of the *cyo* locus as an example is provided in **Figure 4a**. In addition to the cytochrome oxidases, the gene *ygiN* was deleted as well: this gene encodes a probable quinol monooxygenase that uses molecular oxygen to oxidize quinols to quinones^64^. Previous reports suggest that *ygiN* is upregulated in the absence of cytochrome oxidases, resulting in a non-negligible level of oxygen consumption in otherwise fermentative cells^29, 63^. We found this to be true as well and deleted *ygiN* through replacement of the wild-type coding DNA sequence with an ampicillin/carbenicillin resistance cassette (Δ*ygiN*::*amp*).

**Figure 4:**
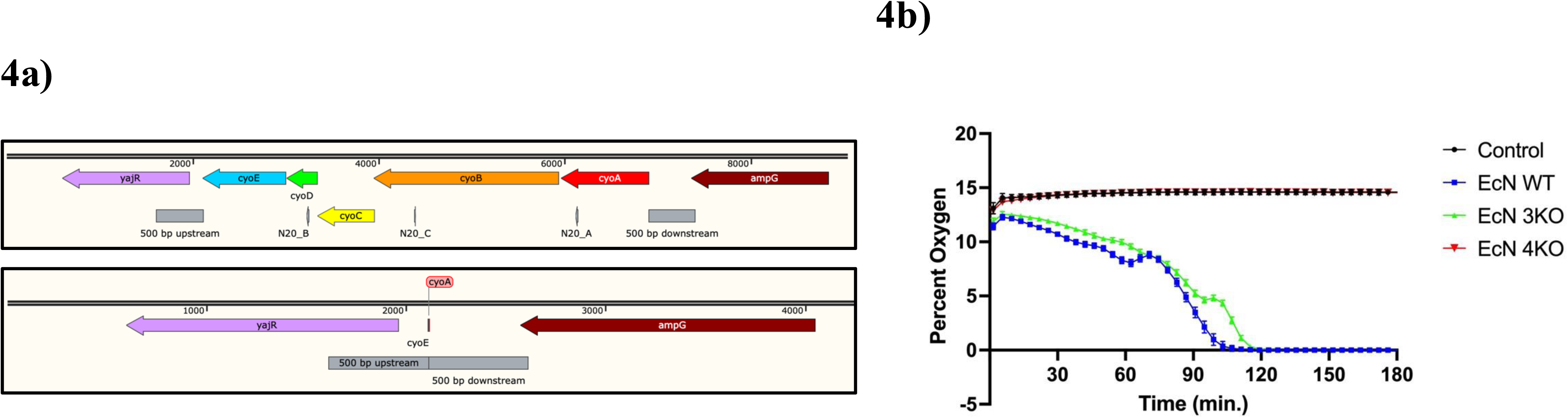
(a) Depiction of gene deletion strategy in *E. coli* Nissle as described in the “Materials and Methods” section. Images from SnapGene. (b) Measurement of extracellular oxygen levels using Oxodish system. Medium was left uninoculated (control) or inoculated with the OD600-normalized strain. Both EcN WT and EcN 3KO are statistically less than both control and EcN 4KO wells (p < 0.05) at all data points, while control and EcN 4KO are not statistically different (p ≥ 0.05). Statistical significance determined using two-way ANOVA. Error bars represent standard deviation.

Although at least one group has observed loss of oxygen utilization in a YgiN-expressing strain, we found that deletion of *ygiN* was necessary in order to completely abolish oxygen consumption^62^. The resulting EcN “quadruple” knockout (Δ*cyo*ABCDE Δ*appCB* Δ*ygiN*::*amp* Δ*cydAB*) was termed “EcN 4KO” and demonstrated poor growth in LB-Miller medium, as previously found for *E. coli* MG1655 with a similar genotype^29, 63^. We observed good growth in BHI, however, and used this medium for growth of EcN 4KO. We also generated “EcN 3KO” as a control, which was identical to EcN 4KO except that it still expressed the cytochrome *bd*-I oxidase Cyd (Δ*cyo*ABCDE Δ*appCB* Δ*ygiN*::*amp*). All strains that were used at the same time as EcN 4KO were grown in BHI to control for any potential effects of bacterial medium on observed phenotype. We then sought to demonstrate that EcN 4KO was deficient in oxygen respiration – to this end, we incubated EcN WT, EcN 3KO, and EcN 4KO in Oxodish plates as done previously to measure extracellular oxygen levels^20^. As shown in **Figure 4b**, EcN WT robustly consumes dissolved oxygen, rendering the Oxodish wells hypoxic within two hours. EcN 3KO, which contains a functional cytochrome oxidase (Cyd), similarly consume oxygen rapidly but with slightly slower kinetics – this could be due to the lower efficiency of the cytochrome *bd*-I oxidase in generating a proton motive force versus the main aerobic cytochrome *bo* oxidase ^65^. In contrast to EcN WT and EcN 3KO, the “quadruple mutant” EcN 4KO shows no oxygen consumption and is not significantly different at any data point versus the uninoculated control wells (p > 0.05 by two-way ANOVA).

We next assessed the potential for the EcN 4KO mutant to stabilize HIF-1α in intestinal epithelial cells. We found that this mutant was incapable of stabilizing HIF-1α in C2BBe1 cells, unlike wild-type EcN and the oxygen respiring EcN 3KO mutant (**Figure 5a**). Use of increased inocula of EcN 4KO did not result in restoration of HIF-1α stabilization, suggesting that the absence of HIF-1α is not due to a growth defect of EcN 4KO but rather due to altered metabolic activities (**Supplemental Figure 1**). Additionally, we observed no induction of HIF-1α target genes by qPCR in C2BBe1 cells treated with EcN 4KO as compared with cells treated with EcN WT or EcN 3KO, further supporting the observation that HIF-1α signaling is not induced by treatment with the EcN 4KO mutant (**Figure 5b**). Interestingly, we did observe a significant increase in *VEGFA* expression in the EcN 4KO-treated cells versus control, untreated cells, despite no significant increase in other HIF-1α target genes assayed. We hypothesize that the increased expression of *VEGFA* in this treatment group could be due to the production of lactate from the EcN 4KO bacteria: previous reports indicate that deletion of all three cytochrome oxidases from *E. coli* results in substantial lactate production during aerobic fermentation, and it has also been shown that lactate induced VEGFA transcription/secretion in a variety of cell types and tissues^29, 63, 66–68^. It is worth noting, however, that the increase in *VEGFA* expression due to EcN 4KO treatment is significantly lower than that of the EcN WT and EcN 3KO treatments, indicating that HIF-1α has a positive contribution towards VEGFA expression in EcN-treated epithelial cells. We also performed luciferase reporter assays using HeLa cells transfected with plasmids expressing either firefly luciferase driven by hypoxia response elements (HREs, derived from the HIF-1α target gene *PGK1* promoter) or *Renilla* luciferase constitutively expressed under a cytomegalovirus (CMV) promoter. Treatment of these co-transfected HeLa cells with 300 μM CoCl_2_, EcN WT, or EcN 3KO led to a significant induction of *Renilla*-normalized firefly luciferase signal relative to untreated, transfected control cells (**Figure 5c**) indicating an increase in HRE-dependent transcriptional activity. Yet, treatment of these same cells with EcN 4KO resulted in no significant increase in *Renilla-*normalized firefly luciferase signal (**Figure 5c**), further demonstrating that EcN 4KO does not drive HIF-1α-dependent gene expression. We also sought to confirm our findings using nontransformed intestinal epithelial cells. We found that, similar to our previous observations, EcN stabilizes HIF-1α in nontransformed hIEC-6 cells in a manner dependent on its ability to respire oxygen (**Figure 6a**). An increase in the initial inoculum of EcN “4KO” (EcN 4KO x4, last lane, **Figure 6a**) did not restore HIF-1α stabilization indicating the importance of oxygen respiration to HIF activation.

**Figure 5:**
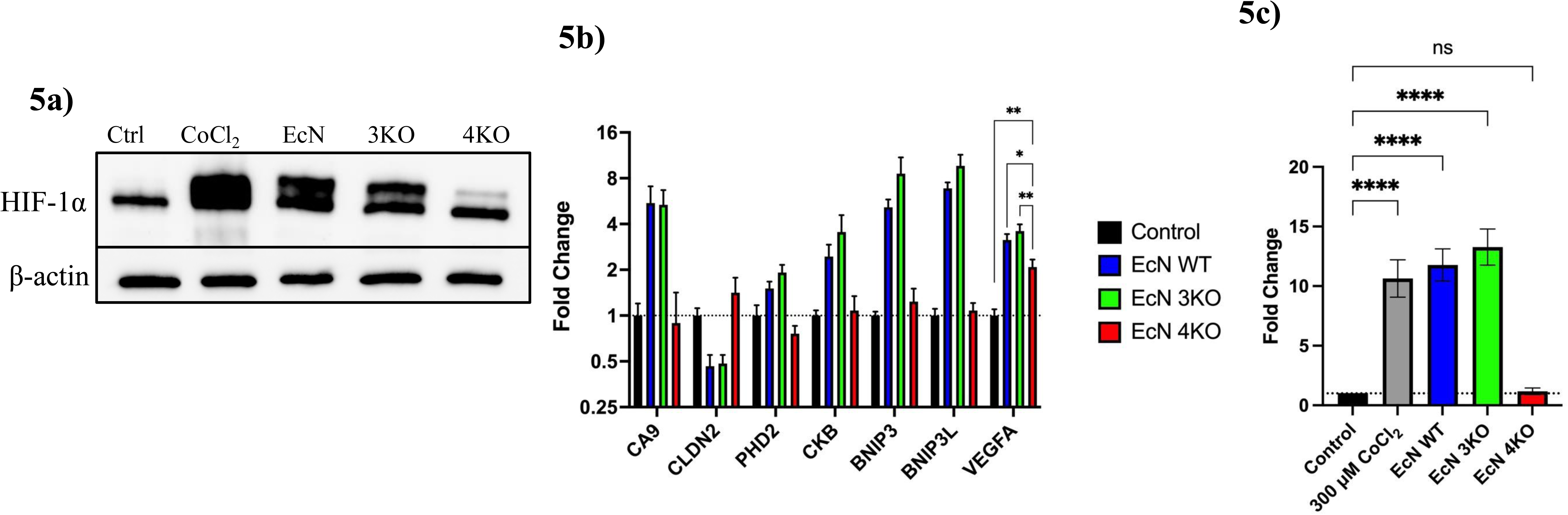
*E. coli* Nissle deficient in aerobic respiration does not stabilize HIF-1α. (a) C2BBe1 cells were treated as indicated for approx. 6 hours, then lysed for analysis by Western blotting. “Ctrl” = untreated (negative) control, “CoCl_2_” = 300 μM cobalt chloride (positive control), EcN = *E. coli* Nissle WT, “3KO” = EcN Δ*cyoABCDE* Δ*appCB* Δ*ygiN*::*amp*, “4KO” = EcN Δ*cyoABCDE* Δ*appCB* Δ*ygiN*::*amp* Δ*cydAB*. (b) Expression of HIF-1α target genes in approx. 8 hr. treated C2BBe1 cells. Data are normalized to β-actin expression and presented a fold-change relative to untreated control cells. All pair-wise comparisons to control are significant (p < 0.05) for EcN WT and EcN 3KO by unpaired t tests with Welch’s correction. In contrast, all pair-wise comparisons to control for EcN 4KO are not significant (p ≥ 0.05) except for *VEGFA*, as indicated. * = p < 0.05 and ** = p < 0.01. (c) Luciferase assay using transfected HeLa cells treated as indicated for approx. 8 hours. Data are presented as firefly luciferase signal normalized to *Renilla* luciferase signal for each sample, then normalized to control (untreated) cells. Statistics calculated pair-wise vs. control using unpaired t tests with Welch’s correction, with **** = p < 0.005 and “ns” = not significant (p ≥ 0.05).

**Figure 6:**
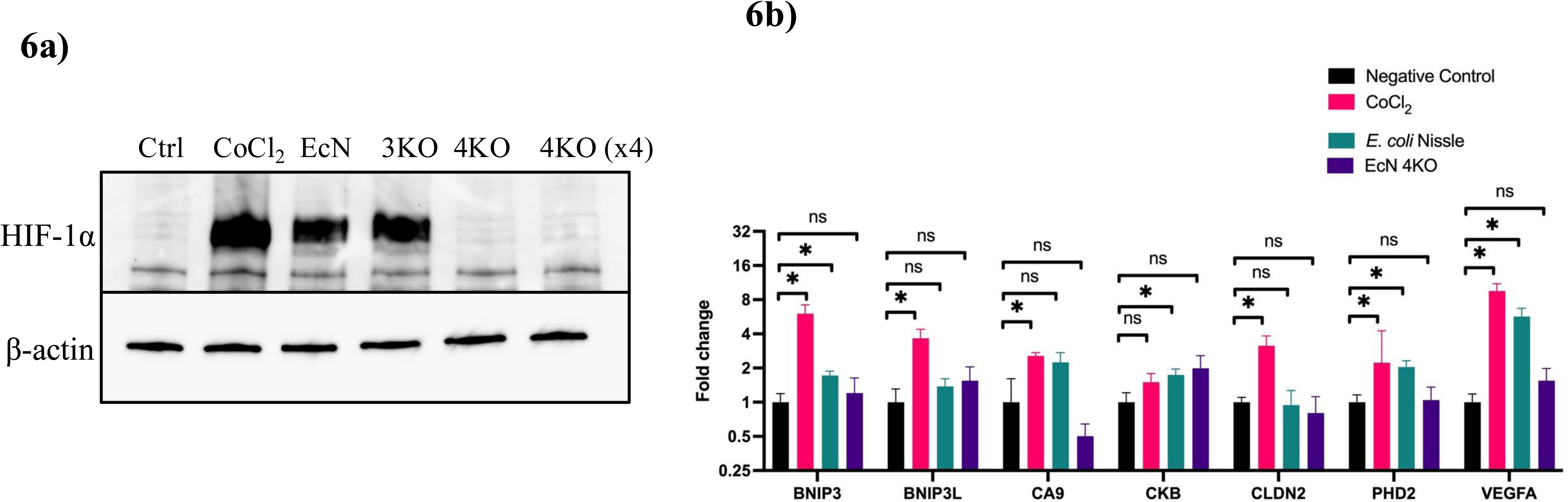
*E. coli* Nissle stabilization of HIF-1α in primary intestinal cells and induction of HIF target genes. (a) hIEC-6 cells were treated with either 300 μM CoCl2 or the indicated strain. “x4” indicates that four times the usual inoculum of EcN 4KO was applied to that sample. (b) HIF target genes were measured in hIEC-6 cells treated with either 300 μM CoCl2 or the indicated strain. Data are normalized to β-actin expression and presented a fold-change relative to untreated control cells. Statistics calculated pair-wise vs. control using unpaired t tests with Welch’s correction. * = p < 0.05 and “ns” = not significant

Additionally, hIEC-6 cells treated with EcN showed induction of HIF target genes similar to our previous findings (**Figure 6b**); this induction was likewise dependent on the ability of EcN to respire oxygen as gene induction was not observed in the EcN 4KO strain. These results, taken together, illustrate that aerobic respiration in *E. coli* Nissle drives HIF-1α stabilization and transcriptional activity in epithelial cells, including upregulation of pro-barrier gene targets, and that abolishing oxygen consumption in EcN is sufficient to nullify these observed phenotypes.

Finally, we sought to confirm our findings using an *in vivo* model of host-microbe interactions. Previous work by our group indicates that intestinal hypoxia is dependent on the gut microbiota, as depleting intestinal bacteria using broad-spectrum antibiotics abolishes pimonidazole (Hypoxyprobe) staining, a marker of tissue hypoxia^36^. Further, we have also previously shown that pimonidazole staining strongly correlates with tissue HIF-1α expression, indicating that detection of pimonidazole can be used as a facsimile for tissue HIF stabilization^69^. To this end, we treated C57BL/6 mice with broad-spectrum antibiotics as done before and assessed pimonidazole staining by in cecal tissue by immunohistochemistry, a commonly used method for measurement of tissue pimonidazole in FFPE samples^36, 70^. Our results show that treatment with antibiotics diminished tissue pimonidazole staining, as indicated by the decreased intensity of brown peroxidase stain in the “+Abx, no EcN” group (**Figure 7**). As a control, the “-Abx” (untreated mice) control tissue was also stained with only the secondary antibody, the absence of brown peroxidase development demonstrating specificity of the anti-pimonidazole primary antibody used. When antibiotic-treated mice were gavaged with EcN, we observed restoration of tissue hypoxia in mice that received wild-type EcN but not those colonized with EcN “4KO”. These findings suggest that wild-type EcN, which rapidly respires O_2_ *in vitro*, is able to restore tissue hypoxia signatures *in vivo* through consumption of luminal oxygen and that the respiration-deficient EcN “4KO” strain lacks this ability.

**Figure 7:**
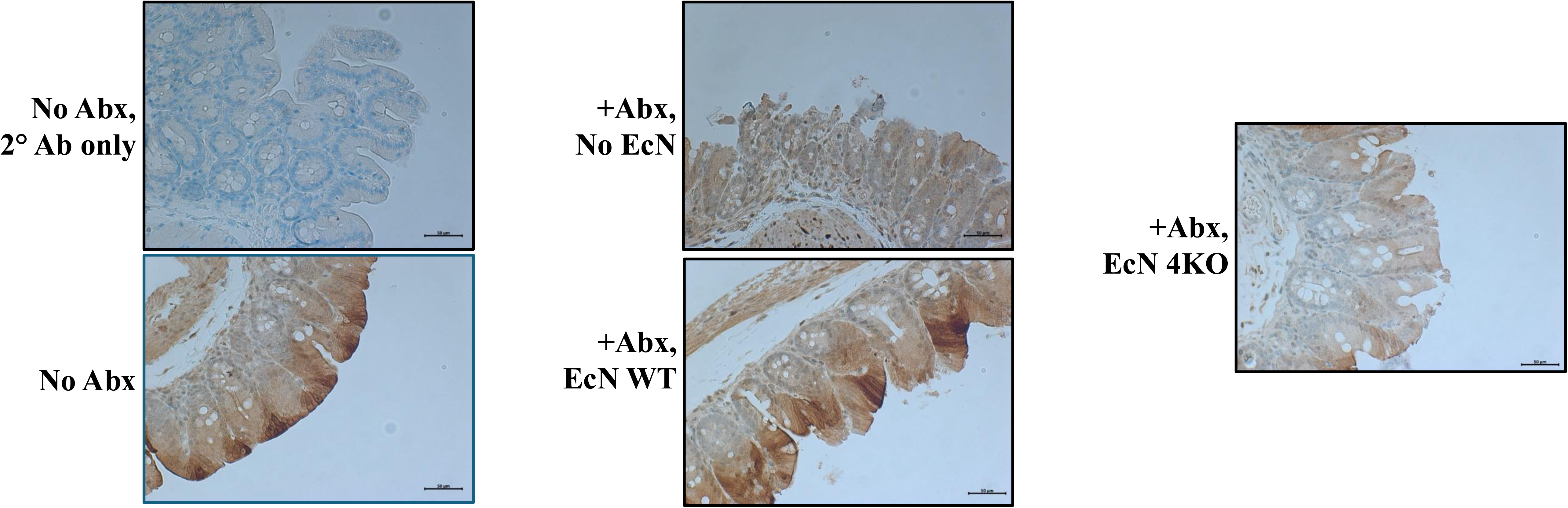
E. coli Nissle promotes tissue hypoxia in vivo. C57BL/6 wild-type mice were treated with broad-spectrum antibiotics (Abx) for three days, then gavaged with either *E. coli* Nissle wild-type (EcN WT) or *E. coli* Nissle Δ*cyoABCDE* Δ*appCB* Δ*ygiN*::*amp* Δ*cydAB* (EcN “4KO”). Mice were treated with pimonidazole (Hypoxyprobe) 1 hour before sacrifice, and cecal tissue was stained using an anti-pimonidazole antibody by immunohistochemistry. Control mice received no antibiotics or antibiotics and no *E. coli*. Representative images shown of two experimental repeats. As a control, a section of cecal tissue from mice that did not receive antibiotics was treated with only secondary antibody to demonstrate specificity of the primary.

## Discussion

Although the mechanisms of intestinal hypoxia have been well-understood for some time, namely β-oxidation of short-chain fatty acids and countercurrent vasculature organization, the full contribution of the microbiota to tissue hypoxia (and, therefore, metabolism and gene expression) is not fully understood^71^. Here, we demonstrate a role for commensal, probiotic *E. coli* in the regulation of intestinal homeostasis through activation of HIF-1α. We show that this regulation occurs through bacterial aerobic respiration, and genetic elimination of oxygen utilization abolishes the observed HIF-1α stabilization and transcriptional activity in epithelial cells. We demonstrate these phenomena in both cancer cell lines and non-transformed intestinal epithelial cells. Finally, we recapitulate our *in vitro* findings by demonstrating that tissue hypoxia can be regulated *in vivo* by *E. coli* Nissle in a manner dependent on its ability to respire oxygen. Our current experimental model based on these findings is summarized in **Figure 8**.

**Figure 8:**
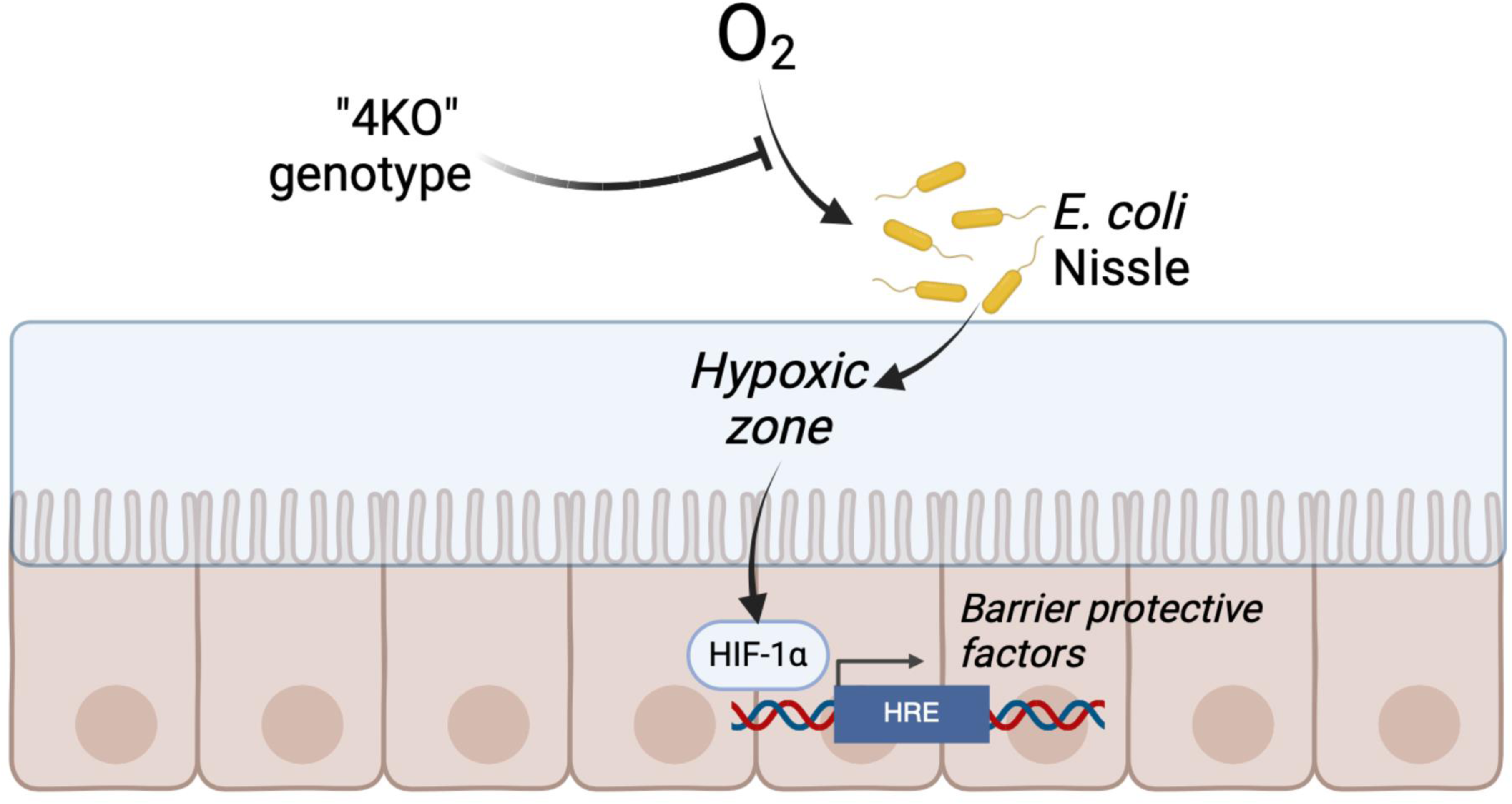
Proposed mechanism for oxygen-dependent regulation of HIF-1α by E. coli Nissle. Oxygen is consumed by E. coli Nissle through aerobic respiration, resulting in a “zone” of hypoxia immediately adjacent to the intestinal epithelium. This results in stabilization of HIF-1α and induction of HIF-1α target genes, which have been previously shown to be protective during intestinal inflammation and to promote gut homeostasis. The loss of bacterial respiration, such as through loss of aerobic respiration in the “4KO” strain, abolishes this “hypoxic zone” and the corresponding HIF-1α stabilization/signaling. Image made using BioRender.

Previous work regarding host-microbe interactions in the gut has demonstrated that the commensal gut microbiota plays an essential role in excluding opportunistically harmful microbes (“pathobionts”) and professionally pathogenic microorganisms (“pathogens”) from establishing themselves in the gut microenvironment. This is accomplished through diverse mechanisms, such as competition for scarce nutrients, secretion of interkingdom signaling metabolites, and regulation of host immunological programs^72^. The loss of “beneficial” microbial species and the proliferation of pathobionts and pathogens, often observed during intestinal inflammation and referred to as “dysbiosis”, is accompanied by numerous genetic and environmental alterations in the gut microenvironment^73^. One major dysbiotic alteration observed during these circumstances is the newfound availability of terminal electron acceptors such as molecular oxygen and nitrate, which promote the growth of facultative anaerobes such as those of the *Enterobacteriaceae* family^74^. This diffusion of oxygen into the intestinal lumen occurs in large part due to a weakening of intestinal barrier function and the loss of bacteria-derived SCFAs, which fuel IEC metabolism through β-oxidation reducing luminal oxygen diffusion^75^.

Interestingly, other groups have observed that oxygen respiration may underlie host-microbe interactions at the intestinal epithelium. Recent work by Litvak et al. demonstrated that competition for oxygen underlies the pathogenesis of *Salmonella* Enteritidis in neonate chicks, and that germ-free animals are protected from infection when colonized with *E. coli* Nissle^76^. This protection was strictly dependent on the ability of EcN to respire oxygen in microaerophilic conditions, as genetic deletion of *cydA* and *appC* resulted in a loss of this protective phenotype. This is in agreement with our findings and those of others, namely that loss of cytochrome oxidases imposes an obligate fermentative phenotype upon *E. coli*^29, 63^. Similarly, blooms of *Candida albicans*, often observed following courses of antibiotics, are dependent on an oxygenated intestinal lumen – the presence of respiring bacteria can combat these fungal expansions in a similar manner as *S.* Enteritidis^77^. Perhaps most intriguing is the reported ability of *E. coli* to scavenge and detoxify host reactive oxygen species (ROS) encountered during intestinal inflammation^78^. Although host-derived nitrates have long been known to be an electron acceptor for various enteric bacteria such as *S.* Typhimurium, recent work by Chanin et al. suggests that the mechanisms for ROS recovery may be widespread amongst enteric bacteria, possibly explaining the well characterized bloom of Proteobacteria observed during intestinal inflammation^78–80^.

Previous work by our group and others has demonstrated that HIF-1α plays an essential role in maintaining intestinal homeostasis. Conditional deletion of this protein in intestinal epithelial cells increased severity of TNBS-induced colitis in a murine model of IBD; likewise, overexpression of HIF-1α through conditional deletion of Vhlh or treatment with PHD inhibitors likewise improves endpoints in animal colitis models^15–17^. Targeting HIF as a route for treatment of intestinal inflammation has long been speculated, though currently no FDA-approved therapeutics for IBD are available that act through this mechanism^81^. The ability of probiotic, noninvasive *E. coli* Nissle to stabilize HIF-1α and potentiate HIF target gene expression suggests that engineering these bacteria to actively consume luminal oxygen may be a viable strategy for future biotherapeutic development. Oral *E. coli* Nissle preparations, including those utilizing engineered strains, have undergone multiple clinical trials in which the treatments were determined to be safe and free from major adverse events^54, 82, 83^. *E. coli* Nissle is genetically tractable, as demonstrated by our work and others, and it is feasible that a strain could be developed that conditionally increases oxygen respiration in the intestinal tract. Such an inducible expression system might be tied to the acidic microenvironment of the inflamed intestine or to the loss of free zinc due to the action of the inflammation-associated protein calmodulin; indeed, expression systems for *E. coli* have already been developed that utilize these environmental signals to regulate gene expression^84, 85^. A strain of *E. coli* Nissle that inducibly “sponges up” luminal oxygen might thus have the multiform effects of inducing HIF signaling (thus, improving intestinal barrier and wound healing), suppressing outgrowth of dysbiotic facultative anaerobes, and permitting re-establishment of SCFA-producing obligate anaerobes.

In summary, our work demonstrates a novel role for probiotic, commensal bacteria in regulating homeostasis of the intestinal epithelium through oxygen respiration and, thus, activation of HIF signaling.

## Data Availability

All data used in preparation of this manuscript is either included herein or is available upon reasonable request to the corresponding author.

## Acknowledgements

This work was supported by funding from the U.S. Department of Veterans Affairs (ASD: 5IK2BX006088; SPC: 5IK6BX006475, 5I01BX002182; IMC: 5IK2BX005710), the National Institute of Diabetes and Digestive and Kidney Diseases (SPC: 2R01DK095491, 5R01DK050189, 2R01DK104713) and the Crohn’s and Colitis Foundation (ASD: 929250).

---

Supplemental Figure 1: “Dose-response” of EcN 4KO. HeLa cells were treated for 6 hours with 300 μM cobalt chloride (“CoCl_2_”) or with the indicated dose of EcN 4KO, then lysed for Western blotting. Control cells (“Ctrl”) were left untreated.

