## Supplementary figures and images for "Regulation of Epithelial HIF by Probiotic *Escherichia coli*"

### Supplemental Figure 1

#### Supplemental Figure 1

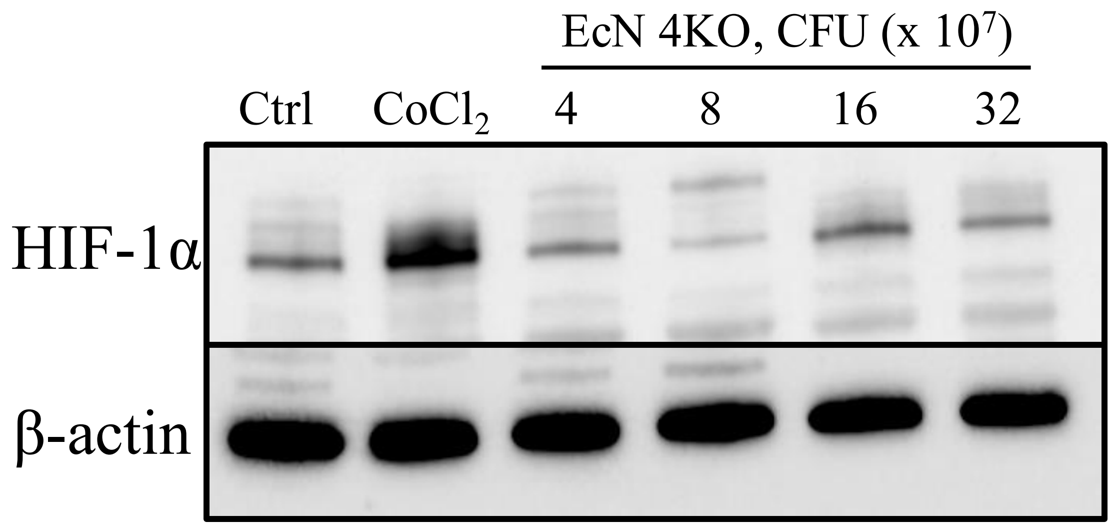
